# Persistence of SARS-CoV-2 neutralizing antibodies longer than 13 months in naturally-infected, captive white-tailed deer (*Odocoileus virginianus*), Texas

**DOI:** 10.1101/2022.07.19.500662

**Authors:** Sarah A. Hamer, Chase Nunez, Christopher M. Roundy, Wendy Tang, Logan Thomas, Jack Richison, Jamie S. Benn, Lisa D. Auckland, Terry Hensley, Walter E. Cook, Alex Pauvolid-Corrêa, Gabriel L. Hamer

## Abstract

After identifying a captive herd of white-tailed deer (WTD) in central Texas with >94% seroprevalence with SARS-CoV-2 neutralizing antibodies in September 2021, we worked retrospectively through archived serum samples of 21 deer and detected seroconversion of all animals between December 2020 and January 2021. We then collected prospective samples to conclude that the duration of persistence of neutralizing antibodies is at least 13 months for 19 (90.5%) of the animals, with two animals converting to seronegative after six and eight months. Antibody titers generally waned over this time frame, but three deer had a temporary 4 to 8-fold increases in PRNT titers over a month after seroconversion; anamnestic response cannot be ruled out.

## Introduction

Approximately 30-40% seroprevalence for SARS-CoV-2 has been reported in wild white-tailed deer (WTD, *Odocoileus virginianus*) across Midwestern states and Texas [1,2]. Experimental studies have corroborated these findings demonstrating that WTD lung cells are permissive to SARS-CoV-2 infection, and that WTD can transmit SARS-CoV-2 vertically and through contact [3,4]. Field investigations suggest sustained horizontal transmission among deer in nature [5,6]. The increasing evidence of multiple spillover events from humans to WTD followed by deer-to-deer transmission includes the detection of several lineages of SARS-CoV-2 in WTD including those that were dominant as well as uncommon in the human population [5], raising concern for novel variants and spillover to humans [7]. Despite the potential of WTD as a natural SARS-CoV-2 reservoir, critical information including the time course of natural infections and persistence of neutralizing antibodies in this species is unknown.

In our previous study, 34 (94.4%) of 36 WTD sampled in September 2021 in one of three captive deer facilities visited in central Texas were seropositive for SARS-CoV-2 [8]. The seroprevalence in this captive herd was more than double that of free-ranging WTD [1,2], suggesting that confined environments may facilitate transmission. We sought to identify the time of initial exposure and determine antibody persistence in this naturally infected population.

Twenty-one does from a captive cervid facility in Texas (19 that tested seropositive for SARS-CoV-2 by PRNT_90_ on September 15 of 2021 and two that tested seronegative) were the subject of the current investigation. All 21 deer were previously enrolled in experimental tick or anthrax vaccine studies involving biweekly to monthly blood collections between November 2020 to July 2021. Retrospective serum samples from five of those time points were used in the current investigation, in addition to the September 2021 collection that sparked the current study. Prospectively, one more blood draw and swab sampling was conducted on March 4th, 2022, following methods previously reported [8]. Animal use was overseen by TAMU’s Institutional Animal Care and Use Committee (2018-0460 CA). Serum was tested for SARS-CoV-2 by plaque reduction neutralization tests (PRNT_90_) using Isolate USAIL1/2020, NR 52381 (BEI Resources, Manassas, VA), and swab samples were tested by RT-qPCR to amplify the SARS-CoV-2 RdRp gene, as previously reported [8,9].

## Results and Discussion

Twenty-one does from a captive cervid facility in central TX had serum tested for SARS-CoV-2 neutralizing antibodies across seven time points from 9 November 2020 – 4 March 2022, with respiratory and rectal swabs tested by RT-qPCR at the last two time points. All does were seronegative in November and December of 2020, and all tested positive by PRNT_90_ in January of 2021, confirming that the time of infection and subsequent seroconversion occurred between 16 December 2020 and 27 January 2021. Except for two animals (Deer-013 and Deer-025) that were seronegative by July and September, most animals (90.5%) had neutralizing antibodies for SARS-CoV-2 detectable from initial seroconversion (January 27, 2021) to the end of the study (March 4, 2022), suggesting that antibodies persisted for at least 13 months (402 days) (Figure 1). Titers generally decayed over time (in some cases with 2-fold temporary increase in titer which alone were interpreted as biologically insignificant), but Deer 10 had a 4-fold increase in titer between January-March 2021, and Deer-006 and 011 had 4-8 fold increases in titers between September 2021 and March 2022.

**Figure 1.**
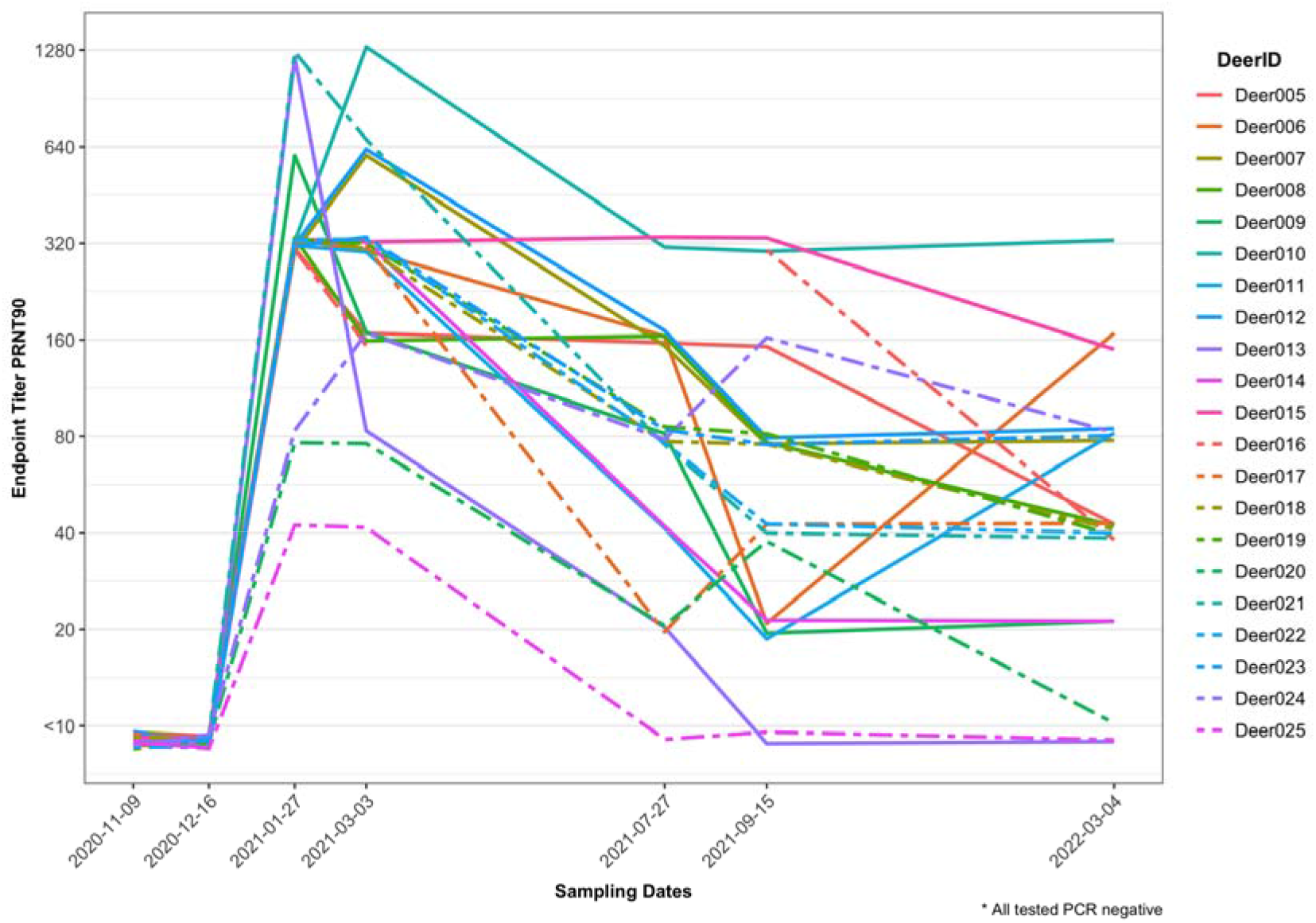
PRNT_90_ endpoint titers for SARS-CoV-2 in 21 white-tailed deer does from central Texas across seven time points over a 16-month period. Deer 006, 010, and 11 showed 4-8 fold increases in titer at least one month after initial seroconversion.

The geometric mean of endpoint PRNT_90_ titers decreased only gradually from 299.6 in January 2021 to 254 in March 2021, with a more rapid decrease thereafter, to 66.5 in by July 2021. The rate of decline then slowed again, to 46.2 in September 2021 and 36.7 in March 2022 (Supplemental Table). The RT-qPCR results from respiratory and rectal swabs of all deer were negative in the September 2021 timepoint as previously reported [8] and were negative in the March 2022 timepoint.

An increasing number of reports suggest that the WTD may be a key species for a sustainable enzootic cycle of transmission of SARS-CoV-2. In this ~16-month longitudinal study, we examined the neutralizing antibody kinetics of a naturally-infected WTD population and found specific neutralizing antibodies for at least 402 days. Because this study was uncontrolled under field conditions using mostly retrospective serum samples, we cannot discount the possibility that multiple SARS-CoV-2 exposures and/or secondary humoral responses occurred. With rare exceptions, we observed a continuous decrease in neutralizing antibody titers over 13 months. Between January of 2021 and March of 2022, the geometric mean of endpoint titers decreased 88% which may support an initial single group of exposure events rather than multiple exposures over time.

All 21 does were initially seropositive on January 27 of 2021, suggesting the time of infection likely occurred between mid-December 2020 and mid-January 2021. In our initial exploratory sampling in September 2021, we reported 94.4% seroprevalence; the only two deer that were seronegative in that study were herein found to be seropositive one or two timepoints prior, suggesting they had also seroconverted with the rest of the population between December 2020 and January 2021. Similarly, in a study of over 280 WTD in Iowa [5], the prevalence of retropharyngeal lymph nodes of WTD that tested positive for viral RNA increased from September through December 2020. Nationwide, there was an increase in human incidence of COVID-19 during November and December of 2020 [10].

Given most deer remained seropositive at the last prospective sampling timepoint, the total length of time that neutralizing antibodies persist in WTD remains unknown. However, our work greatly expands prior estimates based on experimental infection studies which, for obvious reasons, are limited by the empirical dose of infection and restricted time spans [3,4]. Only two (~10%) does had neutralizing antibodies wane below detectable levels, suggesting that depletion of neutralizing antibodies is unlikely to occur in WTD in the first 13 months after natural infection, and that neutralizing antibodies may persist longer. In humans, it has been shown that neutralizing antibodies for SARS-CoV-2 can persist for at least a year [11].

It is noteworthy that the threshold of protective neutralizing antibody endpoint titers for SARS-CoV-2 has not been established for WTD. Therefore, high endpoint titers do not equate to a known protection from reinfection of the same or different viral variants. Experimentally, infected cats produced neutralizing antibodies sufficient to prevent reinfection following a second viral challenge [12]. Further, reinfections caused by different variants have been widely reported in humans [13].

Across the study, most deer had neutralizing antibodies decrease over time, although three deer had a 4 to 8-fold temporary increase titer over the study period. These exceptions include Deer-006 and Deer-011, which both increased between September 2021-March 2022, and Deer-010 which had a 4-fold increase in titer between January and March of 2021. Sequential exposures to SARS-CoV-2 resulting in anamnestic responses have been reported in vaccinated humans [14] and cannot be disregarded for these individuals. Anamnestic responses may obfuscate the rate of neutralizing antibody titer decay.

All the 21 white-tailed does tested here had history of being enrolled in an anthrax vaccine study and a fever tick vaccine study. This was unlikely to influence the SARS-CoV-2 antibody findings given many WTD from another cervid facility were enrolled in the same anthrax vaccine study and showed no neutralizing antibodies for SARS-CoV-2 using the same assay [8]. The PRNT is considered as gold standard for serological testing for coronaviruses [15] and we used a highly conservative criteria of seropositivity (90% neutralization) aiming to mitigate the detection of non-specific heterologous reactions.

Given the high densities of deer in nature, which may be exacerbated in captive cervid settings, these data suggest WTD have the unique potential to act as amplifying hosts of SARS-CoV-2 in nature. The findings presented on the humoral immune response of WTD naturally infected by SARS-CoV-2 strengthen our understanding of natural infection kinetics and raise important questions on whether SARS-CoV-2 neutralizing antibodies can prevent a secondary infection and ultimately the spread of SARS-CoV-2 by infected WTD.

## Acknowledgements

Texas A&M AgriLife Research provided funding. The following reagent was deposited by the Centers for Disease Control and Prevention and obtained through BEI Resources, NIAID, NIH: SARS-Related Coronavirus 2, Isolate USAIL1/2020, NR 52381. Project data are available at the OakTrust digital repository: https://hdl.handle.net/1969.1/196210

No potential competing interest was reported by the authors.

