## Supplemental Table 1 for "Persistence of SARS-CoV-2 neutralizing antibodies longer than 13 months in naturally-infected, captive white-tailed deer (*Odocoileus virginianus*), Texas"

Supplemental Table. PRNT_90_ endpoint titers for SARS-CoV-2 across 16 months in captive white-tailed deer from central Texas, 2020-2022.

| ID | 11/9/20 | 12/16/20 | 1/27/21 | 3/3/21 | 7/27/21 | 9/15/21* | 3/4/2022* |
| --- | --- | --- | --- | --- | --- | --- | --- |
| Deer-005 | <10 | <10 | 320 | 160 | 160 | 160 | 40 |
| Deer-006 | <10 | <10 | 320 | 320 | 160 | 20 | 160 |
| Deer-007 | <10 | <10 | 320 | 640 | 160 | 80 | 80 |
| Deer-008 | <10 | <10 | 320 | 160 | 160 | 80 | 40 |
| Deer-009 | <10 | <10 | 640 | 160 | 80 | 20 | 20 |
| Deer-010 | <10 | <10 | 320 | 1,280 | 320 | 320 | 320 |
| Deer-011 | <10 | <10 | 320 | 320 | 40 | 20 | 80 |
| Deer-012 | <10 | <10 | 320 | 640 | 160 | 80 | 80 |
| Deer-013 | <10 | <10 | 1,280 | 80 | 20 | <10 | <10 |
| Deer-014 | <10 | <10 | 320 | 320 | 40 | 20 | 20 |
| Deer-015 | <10 | <10 | 320 | 320 | 320 | 320 | 160 |
| Deer-016 | <10 | <10 | 320 | 160 | NT | 320 | 40 |
| Deer-017 | <10 | <10 | 320 | 320 | 20 | 40 | 40 |
| Deer-018 | <10 | <10 | 320 | 320 | 80 | 80 | 40 |
| Deer-019 | <10 | <10 | 320 | 320 | 80 | 80 | 40 |
| Deer-020 | <10 | <10 | 80 | 80 | 20 | 40 | 10 |
| Deer-021 | <10 | <10 | 1,280 | 640 | 80 | 40 | 40 |
| Deer-022 | <10 | <10 | 320 | 320 | 80 | 40 | 40 |
| Deer-023 | <10 | <10 | 320 | 320 | 80 | 80 | 80 |
| Deer-024 | <10 | <10 | 80 | 160 | 80 | 160 | 80 |
| Deer-025 | <10 | <10 | 40 | 40 | <10 | <10 | <10 |
| Geometric mean | <10 | <10 | 299.56 | 253.98 | 66.52 | 46.18 | 36.65 |

NT - Not tested; <10 replaced by 1 for geometric mean calculation

*All deer at these two time points had oral and rectal swabs tested for SARS-CoV-2 by RT-QPCR with negative results.
